# Exploring new roles for actin upon LTP induction in dendritic spines

**DOI:** 10.1101/2020.11.14.382663

**Authors:** Mayte Bonilla-Quintana, Florentin Wörgötter

## Abstract

Dendritic spines, small protrusions of the dendrites, enlarge upon LTP induction, linking morphological and functional properties. Although the role of actin in spine enlargement has been well studied, little is known about its relationship with mechanical membrane properties, such as membrane tension, which is involved in many cell processes, like exocytosis. Here, we use a 3D model of the dendritic spine to investigate how polymerization of actin filaments can effectively elevate the membrane tension to trigger exocytosis in a domain close to the tip of the spine. Moreover, we show that the same pool of actin promotes full membrane fusion after exocytosis and spine stabilization.

## Introduction

Dendritic spines are small protrusions from the dendrites where the postsynaptic part of most excitatory synapses are located. At the tip of the spines and opposite to the presynaptic bouton, there is a specialized substructure called the postsynaptic density (PSD) where receptors and signalling molecules are localized. The AMPA-type glutamate receptors (AMPARs) in the PSD are activated when the presynaptic neuron releases glutamate. The spine depolarizes when these receptors are strongly enough activated, allowing calcium entry via N-methyl-D-aspartate receptors (NMDARs). This triggers an increase in spine size that is associated with an increase in AMPA-receptor-mediated currents and depends on NMDARs, calmodulin, and actin polymerization [1]. Such an increase in AMPA-receptor-mediated current is due to an increment of AM-PARs at the synapse by exocytosis or lateral movement [2, 3]. Moreover, it enhances the signal transmission between two neurons, thereby strengthening their synapse in a process called long-term potentiation (LTP). These activity-dependent structural modifications of dendritic spines are thought to be a cellular basis for learning and memory [1].

Spines experience an increase in their volume of ≈ 200%, which happens within five minutes after stimulation [1,4,5]. In some spines, volume increments of ≈ 50% have been observed after 100 minutes of initial stimulation [1], indicating that the initial increase of spine volume after stimulation is transient and that spine volume stabilizes at a smaller value, that is still larger than the initial value. This volume change is possible due to actin, the principal cytoskeleton component in the spine [4]. Actin is a globular protein (G-actin) that assembles into filaments (F-actin), which are polar structures that undergo a continuous treadmilling process, where G-actin with bound ATP is polymerized at the barbed (+) end of the filament. Furthermore, bound ATP hydrolyzes to ADP promoting depolymerization of G-actin at the pointed (-) end of the filament. Actin binding proteins (ABPs) promote polymerization and depolymerization, as well as branching and severing of F-actin.

Within spines, there is a dynamic equilibrium between G- and F-actin [6]. Moreover, actin filaments are organized in two different pools: a dynamic pool localized at the tip of the spine with fast treadmilling velocity, and a static pool with slow treadmilling velocity at the base of the spine head [7]. After activity induction, a third pool, called the “enlargement” pool, forms [7]. Importantly, actin serves as the main anchoring site for many postsynaptic proteins including NMDA and AMPA receptors [8]. Interestingly, the PSD size correlates with spine size and the PSD grows in spines that show persistent enlargement [9].

Upon LTP, the structural changes in spines can be divided into three temporal phases [5]. During the first phase (1-7 min), spines rapidly expand due to an increase in the concentration of ABPs, such as Arp2/3 and Aip1, that modify F-actin through severing, branching and capping. At the same time, the pool of ABPs, which stabilizes F-actin, is transiently depleted. Thus, during this phase there is an increase in actin polymerization at the barbed ends that elongates F-actin, promoted by F-actin severing, resulting in a profound remodelling of the spine. During the second phase (7-60 min) ABPs that modify F-actin and ABPs that promote F-actin stabilization return to their basal levels, thereby stabilizing the F-actin cytoskeleton. In the third phase (>60 min), the concentration of some PSD proteins within the spine increases. These proteins are translocated to the PSD, which results in PSD enlargement, thereby, recovering the relation between spine and PSD size. Note, however, that some studies also show that different PSD proteins experience an almost immediate increase in their concentration upon LTP [9].

Actin polymerization at the barbed ends near the membrane generates a force that pushes the membrane forward [10] and raises the membrane tension locally [11], which is involved in many cell processes. For example, it controls cell motility by producing the signal that coordinates membrane trafficking (via exocytosis), actomyosin contraction, and plasma membrane area change [12]. Moreover, the membrane tension generated by F-actin polymerization facilitates shrinking of the fusion-generated Ω-profiles during exocytotic events [13]. A recent theoretical model shows that membrane tension is important for maintenance of different spines shapes [14]. However, to our knowledge, the effects of spine enlargement after induction of LTP on membrane tension and its consequences in cellular processes, such as exocytosis, have been neglected both experimentally and theoretically.

Here, we use a theoretical 3D model to investigate how changes in spine shape affect membrane tension. Assuming that an increase in membrane tension acts as a mechanical signal that triggers exocytosis, we examine how actin polymerization induces such an increase by simulating different scenarios. Moreover, we study how actin facilitates membrane fusion during exocytosis, and size stabilization. We find that a discrete F-actin polymerization focus located near the PSD increases membrane tension most efficiently and aids membrane fusion and spine stabilization, suggesting that these events co-localize and result from the same actin pool.

## Results

We used a theoretical model, based on [15], in which asymmetric spine shape changes result from an imbalance between a force generated by actin polymerization, that pushes the membrane forward, and the membrane force that counteracts these deformations. To allow for a better measure of the membrane tension, we used a simplified description of the actin dynamics. Moreover, we only considered the actin that is polymerizing close to the spine membrane and, thus, affects its shape.

### Distribution of F-actin polymerization affects the force generated by membrane tension

In the absence of external forces, our modeled spine membrane reaches a resting shape that minimizes the membrane energy (*ε*_*mem*_ in Eq. (4), black shape in Fig. 1A). After obtaining the resting shape for a spine, considering experimental data of spine size and PSD surface area in [9, 16] (see Methods for details), we simulated LTP induction by including a force generated by rapid and persistent F-actin polymerization [6]. This corresponds to the first phase of the structural changes in the spine triggered by LTP induction where actin filaments are continually assembled and disassembled [4]. In our model, this is simulated by a continuous force **F**_*actin*_, given by Eq.(2), that pushes the membrane forward and is inversely proportional to the distance between the starting (nucleation) location of the F-actin polymerization focus **f** ^*i*^ and the spine membrane. Thus, the actin polymerization force is greater near the nucleation location and diminishes as the spine expands. This way, we considered F-actin that branches laterally and has a fast treadmilling velocity.

**Figure 1:**
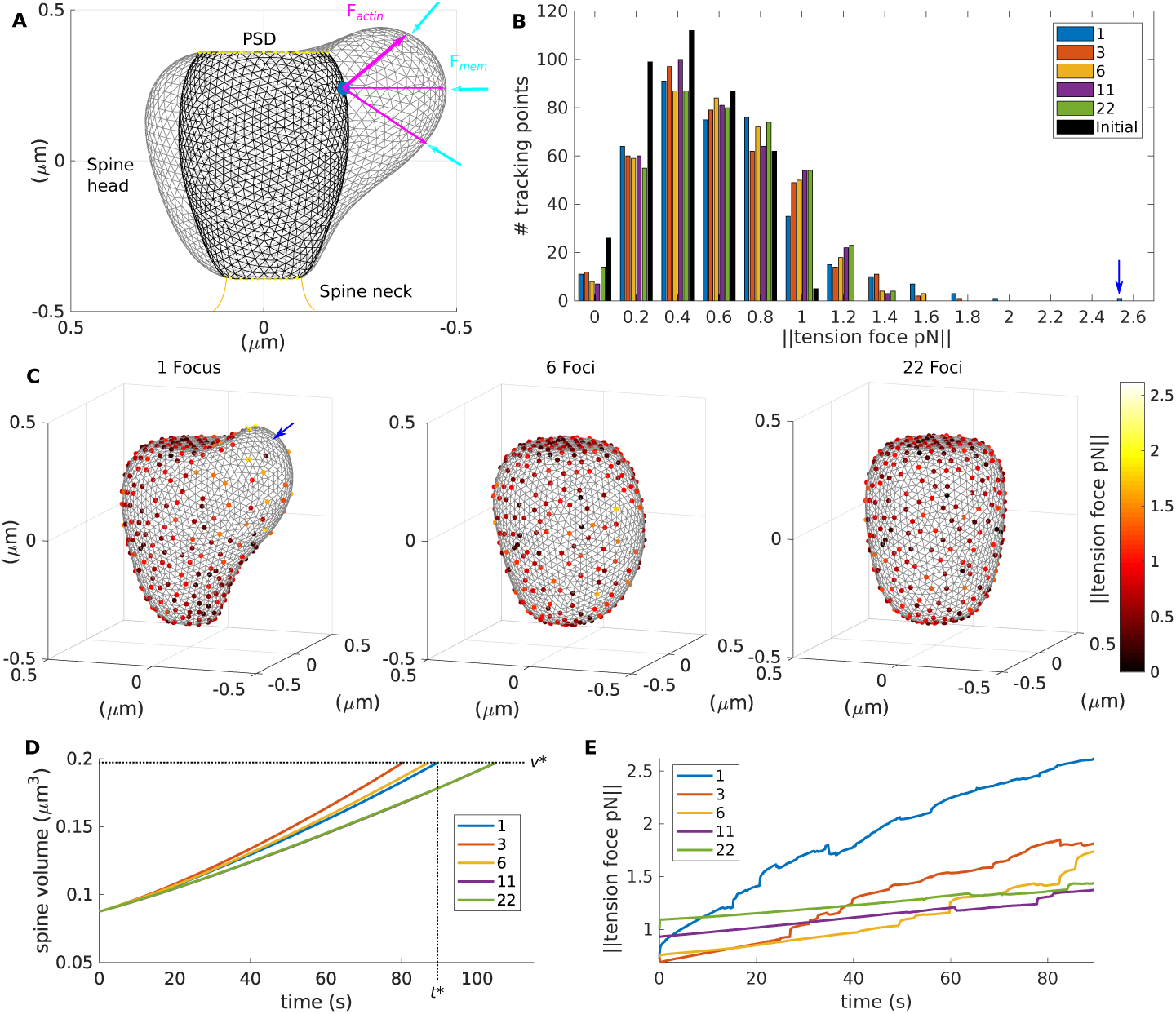
Spine enlargement upon LTP for different number of F-actin polymerization foci. **A**, spine resting shape (black), and spine shape after LTP induction (gray). The membrane deformation results from an imbalance between a force exerted by F-actin polymerization (**F**_*actin*_, magenta arrows) and a membrane force (**F**_*mem*_, cyan arrows). The blue dot signals the location of the F-actin polymerization focus. This plot shows the *y − z* axis for *x* = 0. **B**, histogram of the distribution of the force generated by membrane tension measured at the tracking points when the spines reach a volume equal to *v*^*^. Colors denote the number of polymerization foci distributed in the spine for each simulation. The distribution of the membrane tension for the resting shape, used at the beginning of the simulations, is shown in black. Blue arrow signals maximum tension. **C**, spines shapes (gray) at time *t*^*^ with different number of F-actin polymerization foci. Dots are the tracking points color-coded for the force generated by the membrane tension. **D**, spine volume evolution over time, color-coded by the number of polymerization foci. Note that the traces for 11 and 22 foci are very similar. Dotted black lines denote *t*^*^ and *v*^*^. **E**, evolution of the membrane tension for the tracking point with maximum membrane tension when the spine reaches a volume *v*^*^.

When the membrane deforms due to the actin polymerization force, its energy increases and generates a response force **F**_*mem*_ (Eq. (3)). Figure 1A shows the resulting spine shape in gray. In our simulations, we tracked **F**_*mem*_ and each of its components at different locations along the spine membrane, in particular the work needed to increase the surface area (dots in Fig. 1C). This way, we captured the membrane force generated by tension (tension force) and studied its spatial distribution and evolution over time and whether it depends on the number of actin foci. This method to approximate the membrane tension force resembles experiments that use optical tweezers to calculate tether force [12] in the sense that both obtain the force generated by the displacement of a reference point. In [12] the membrane tension is proportional to the square of the tether force, but here we did not attempt to calculate this relationship and, instead, reported the tension force derived from Eq. (3). During these simulations, the PSD and neck size remained unchanged (yellow dots in Fig. 1A), as observed in experiments within this temporal interval [4, 5, 9].

First, we investigated spine expansion and membrane tension force under different scenarios of F-actin polymerization locations upon LTP induction. To allow comparison between these scenarios, **F**_*actin*_ (Eq. (2)) was multiplied by a constant *ϕ*, that is inversely proportional to the number of polymerization foci. Thus, the force generated by actin polymerization was distributed among the foci.

We found that when actin polymerization accumulates in one location near to the PSD, the membrane tension force increased 2.5 fold at that location at time *t*^*^ = 89.5 seconds. Thereupon, the spine volume is *v*^*^ = 0.1970 m*μ*^3^, growing by ≈ 125%. Spines with more F-actin polymerization foci also increased their volume to *v*^*^, but at different times (Fig. 1D). Because **F**_*actin*_ distributes among the polymerization foci, the time difference to reach the same volume is related to the foci position within the spine. When there are enough foci to be evenly distributed (11 or 22), the spine volume increases similarly, as shown by the purple and green traces in Figure 1D. If there a few foci, two of them could be close enough to cooperate, resulting in a fast enlargement, as in the case with 3 foci. This highlights the complex interaction between **F**_*mem*_ and **F**_*actin*_ in our model that emerges from the spine morphology.

Figure 1C shows spines at *t*^*^ for different number of foci. Note that spines with more foci increased their volume and the force generated by the membrane tension more isotropic. The histogram of this force, measured by the tracking points when they reach a volume of *v*^*^ (Fig. 1B), reveals that only in the spine with one focus it is greater than 2 pN. Moreover, Figure 1E shows that the evolution of the membrane tension force over time, measured at the tracking point with the maximum tension force value at *v*^*^, is lower for spines with more than one focus.

When studying the evolution of the distribution of the tension force among the tracking points, we observed that at the start of the simulation 60.61% of the tracking points measured a force less than 0.6 pN (black bars in Fig. 1B). Different from this, after *t*^*^ only 42.46% of the points in the spine with one focus measured less than 0.6 pN (blue bars in Fig. 1B). Thus, the tension force increases at most locations and has a total increase from 210.6736 pN, at the start of the simulation, to 280.6355 pN at *t*^*^. Spines with more foci also increase the measured tension force at the tracking locations, but the resulting distribution resembles a normal distribution whilst the distribution of the spine with one focus is skewed. Moreover, the sum of the tension force measured at the spines with more than one focus, when they reached a volume of *v*^*^, is less of that of the spine with one focus (272.71 pN, 275.1510 pN, 274.6277 pN, 278.8661 pN for spines with 3, 6, 11 and 22 foci, respectively).

We conclude that after LTP induction, the force generated by membrane tension shows a bigger increase when F-actin polymerization concentrates in a specific location than when it is evenly distributed over the spine. Importantly, this is not an effect of the dependency of the number of foci on the parameter *ϕ* of Eq. (2), as shown in Supplememtary Figure S1, which shows the evolution of a spine with one or 22 foci with the same value of *ϕ*. Since membrane tension serves as a mechanical signal for exocytosis [12] and the force generated by the membrane tension is proportional to the membrane tension, these results are in line with experimental observations of a defined exocytic domain in the spine [17].

### Position of the F-actin polymerization focus affects membrane tension

Next, we investigated whether the location of the polymerization focus affects the force generated by membrane tension. We compared the spine shape resulting from the polymerization focus near the PSD (Fig. 1A) with those having the focus in the middle of the spine head or near the spine neck. Figure 2A shows that the spine shapes, when they reached a volume of *v*^*^, are different, albeit having the same volume. Spines with the polymerization focus far from the PSD reached this volume faster (Fig. 2C). However, Figure 2B shows that, when the spines reach a volume of *v*^*^, the maximum tension force, measured by the tracking points, is higher for the spine with the focus near to the PSD. Moreover, the sum of these forces along the tracking points is also higher in such a spine (280.6355 pN, compared to 275.1624 pN and 278.7298 pN in the spine with the focus in the middle and near the neck, respectively). Likewise, the evolution of the tension force, measured at the tracking points corresponding to the highest tension force in the spines with volume *v*^*^, shows higher values for the spine with the polymerization focus near to the PSD (Fig. 2D).

**Figure 2:**
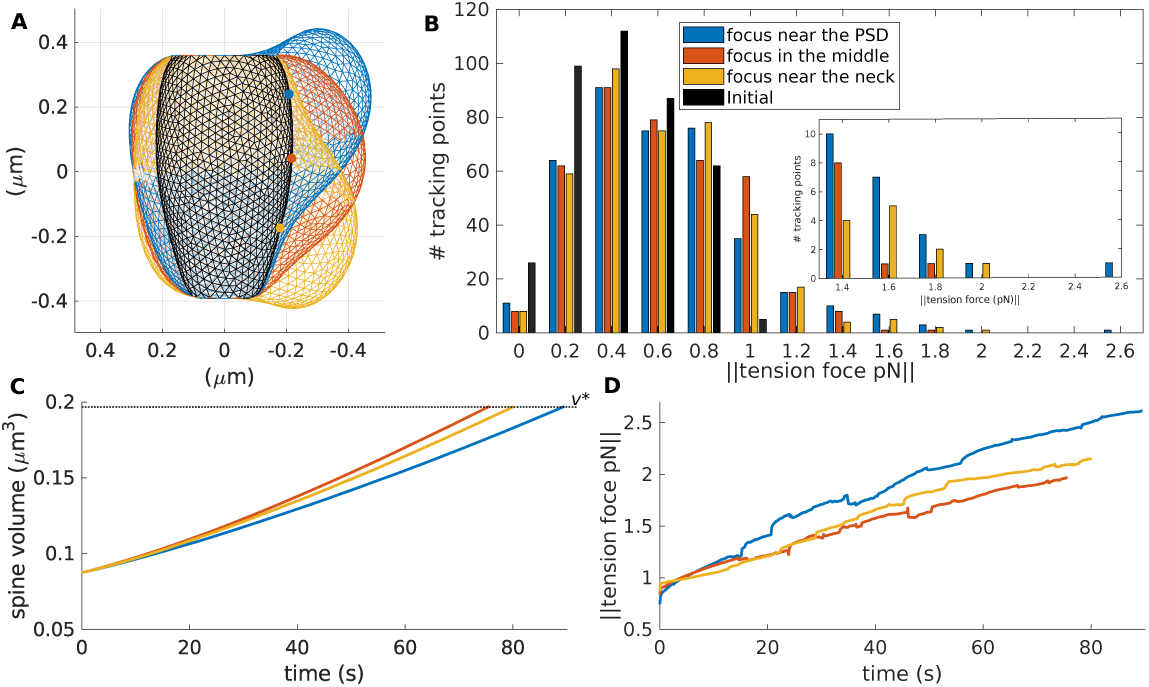
Spine enlargement upon LTP for different locations of the F-actin polymerization focus. **A**, spine shapes for different locations of the F-actin polymerization focus (dots) when they reached a volume of *v*^*^, color-coded as in B. This plot shows the *y − z* axis, for *x* = 0. **B**, histogram of the distribution of the force generated by membrane tension measured at the tracking points when the spines reached a volume of *v*^*^ (this occur at different times, see C). **C**, spine volume evolution over time, color-coded by the number of polymerization foci. Dotted black line denotes *v*^*^. **D**, the evolution of the membrane tension for the tracking point with maximum membrane tension when the spine reaches a volume of *v*^*^.

Note that when the polymerization focus is near the PSD, the membrane stretches more, due to the immobility of the PSD. Although the neck was also fixed, the effect on the force generated by the membrane tension is lower, because the neck surface area is smaller. Taking everything together, the force generated by the membrane tension reaches a higher value when the F-actin polymerization focus is near the PSD, indicating that the exocytic domeain locates therein, as observed in [17].

### F-actin polymerization promotes membrane fusion upon exocytosis

We have shown in our model that when F-actin polymerization concentrates at a certain location near the PSD, it rapidly elevates the force generated by the membrane tension, which can serve as a signal for exocytosis [12]. In the spine, exocytosis is important for AMPARs and membrane trafficking [2, 17, 18]. To achieve membrane fusion and cargo release, vesicles dock to the membrane and form an Ω-shaped structure [13] that has to shrink and merge with the membrane. In our model, shape deformations, like the Ω-profile, generate a response force in the membrane **F**_*mem*_ that restores it to the resting shape. However, Wen *et al*. [13] show that F-actin polymerization mediates Ω-profile merging. Therefore, we investigated two scenarios for Ω-profile shrinking, namely, shrinking resulting only from **F**_*mem*_ dynamics and shrinking aided by F-actin polymerization.

In the first scenario, we assumed that, when exocytosis starts, myosin contracts and pulls the F-actin away from the membrane, as observed in [12]. Consequently, the force generated by actin polymerization ceases. Experimental data shows that this is possible in spines, since myosin II activates upon LTP induction and is required for stabilization of synaptic plasticity [19]. Therefore, we set **F**_*actin*_ = **0** and observed that the Ω-profile depicted in Figure 3A (highlighted in magenta) shrinks fully within 30 seconds (see yellow shapes in Fig. 3D), decreasing spine volume and surface area (Figs. 3B-C). However, exocytosis provides membrane to the spine for enlargement upon LTP [17, 18]. Therefore, there must be an increase in surface area of the spine membrane.

**Figure 3:**
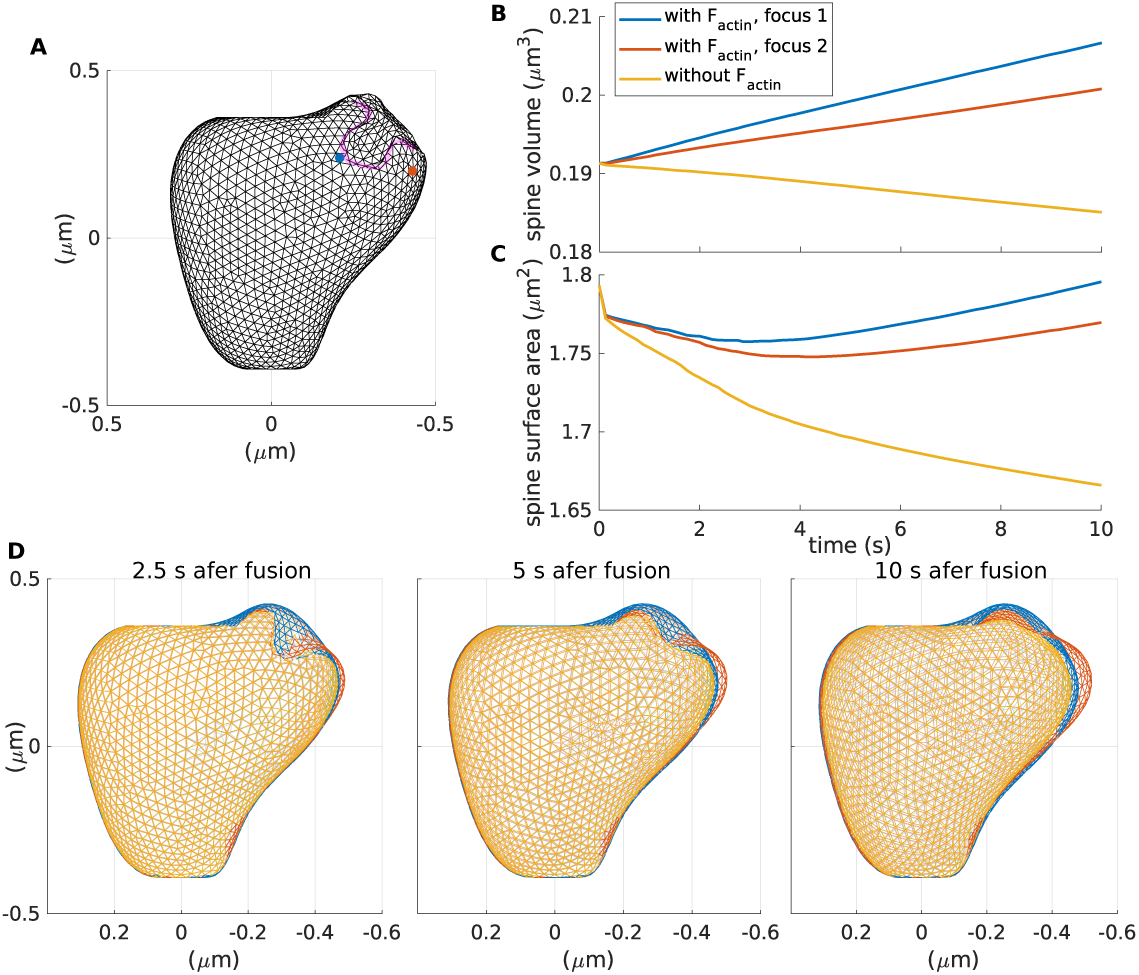
Exocytosis with and without the aid of F-actin polymerization. **A**, dendritic spine membrane after fusion with a recycling endosome. The invagination, highlighted in magenta, is the Ω-profile formed after the fusion event. The blue dot represents the F-actin polymerization focus. **B**, spine volume evolution over time, color-coded depending whether the Ω-profile merging is aided by **F**_*actin*_ or not. **C**, spine area surface evolution over time, color-coded as in B. **D**, snapshots taken at different times after initiation of spine membrane fusion with the recycling endosome, color-coded as in B.

Consequently, we considered a second scenario where actin filaments remain close to the membrane and polymerize. Hence, they generate a polymerization force (i.e. **F**_*actin*_ ≠ **0**) from a focus at the tip of the Ω-profile (blue dot in Fig. 1A). Within 10 seconds, the spine membrane surface area returned to its initial value (Fig. 3C), indicating a possible full merge of the Ω-profile, which supplies membrane. The initial decrease in membrane surface area is transient and could be due the distribution of F-actin. Note that during the simulation, the spine volume increases (Fig. 3B) and the Ω-profile shrinks completely. To test whether the location of the actin focus affects the reduction of the Ω-profile, we ran a simulation with the polymerization focus located next it (orange dot in Fig. 3A). In this case, the volume and surface area of the spine increase (Fig. 3B-C), and the Ω-profile shrinks (Fig. 3D). However, the increase is less than for the previous case and the spine membrane is slightly pushed aside.

These results suggest that actin polymerization is needed for membrane insertion resulting from the full merge of the Ω-profile formed upon exocytosis. Moreover, they indicate that the pool of F-actin that polymerizes and elevates the membrane tension to trigger exocytosis is involved in promoting Ω-profile shrinkage and the full fusion of the vesicle with the spine membrane. Hence, we speculated that there is a pool of F-actin that translocates to a focus near the PSD upon LTP and pushes the membrane forward, which elevates the force generated by the membrane and triggers exocytosis (see Fig. 4). Then, this same pool promotes the shrinkage of the Ω-profile generated when the vesicle docks to the membrane, and hence, full fusion.

**Figure 4:**
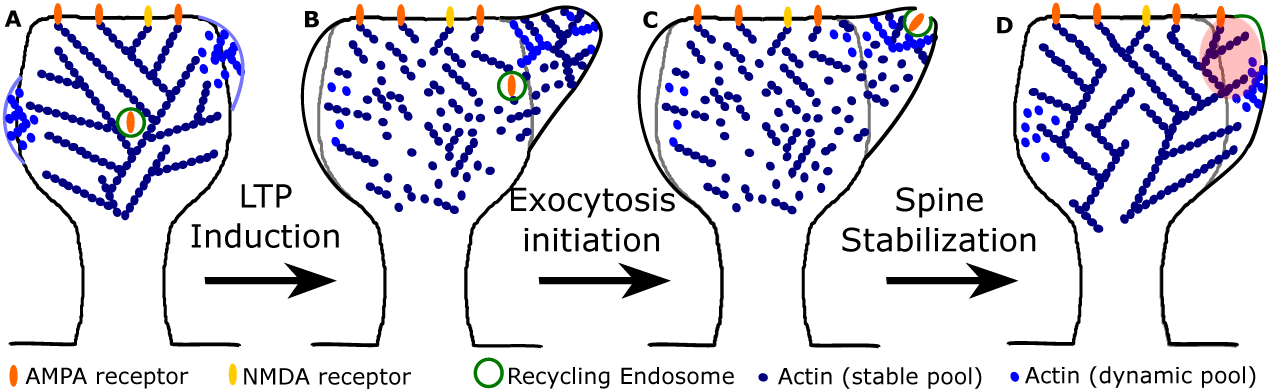
Actin re-organization upon LTP. **A**, before LTP induction, actin distributes in a stable and dynamic pool (dark and light blue dots, respectively). Spontaneous shape fluctuations (light blue lines) result from polymerization of the dynamic pool that organizes in distinct foci [15, 20, 21]. Note that only actin polymerizing close to the membrane can push the membrane forward. **B**, 1 ≈ min after LTP induction, actin rapidly assembles and disassembles. Actin polymerizes at a single location near to the PSD, elevating the force generated by the membrane tension, which triggers exocytosis of the recycling endosome. The initial spine shape is in gray. **C**, actin polymerization promotes full fusion of the Ω- profile formed after the docking of the recycling endosome with the spine membrane. **D**, after completing exocytosis, the spine stabilizes. There is an increase in the AMPARs and spine size. Also, the membrane from the recycling endosome (in green) is merged with the spine membrane. Note that the spine enlargement occurs where the polymerization focus of A was located (highlighted in red).

### Role of the F-actin polymerization focus for spine stabilization

Next, we studied whether the F-actin pool, responsible for triggering and completing the exocytosis events, also accounts for spine stabilization. We hypothesized that after exocytosis, F-actin no longer polymerizes at a fast rate close to the spine membrane. This could be due to the contraction of F-actin promoted by myosin [12] or due to changes in ABPs, namely, an increase in proteins that promote F-actin stabilization and the switch in cofilin function: from severing F-actin to the forming of stable filaments [5].

In our simulations, we increased the radius of the PSD at a rate of Δ_*PSD*_ each time-step, and stabilized the F-actin, corresponding to spine enlargement within this radius, by fixing it at a height of *h*_*PSD*_ (see Methods). Figures 5A-B show the increase in the PSD surface area and the evolution of the spine volume from an initially expanded shape (similar to that in gray in Fig. 1A) for different values of Δ_*PSD*_. Note that larger values of Δ_*PSD*_ reach a higher maximum PSD surface area faster. However, eventually, the PSD surface area settles to a lower value that depends on Δ_*PSD*_. Here, the fluctuations in the evolution are an artifact of the mesh approximation to the membrane. We speculate that the decrease on the PSD area is a consequence of spine shrinkage due to the membrane force that moves the vertices, corresponding to the enlarged PSD, away from *h*_*PSD*_.

**Figure 5:**
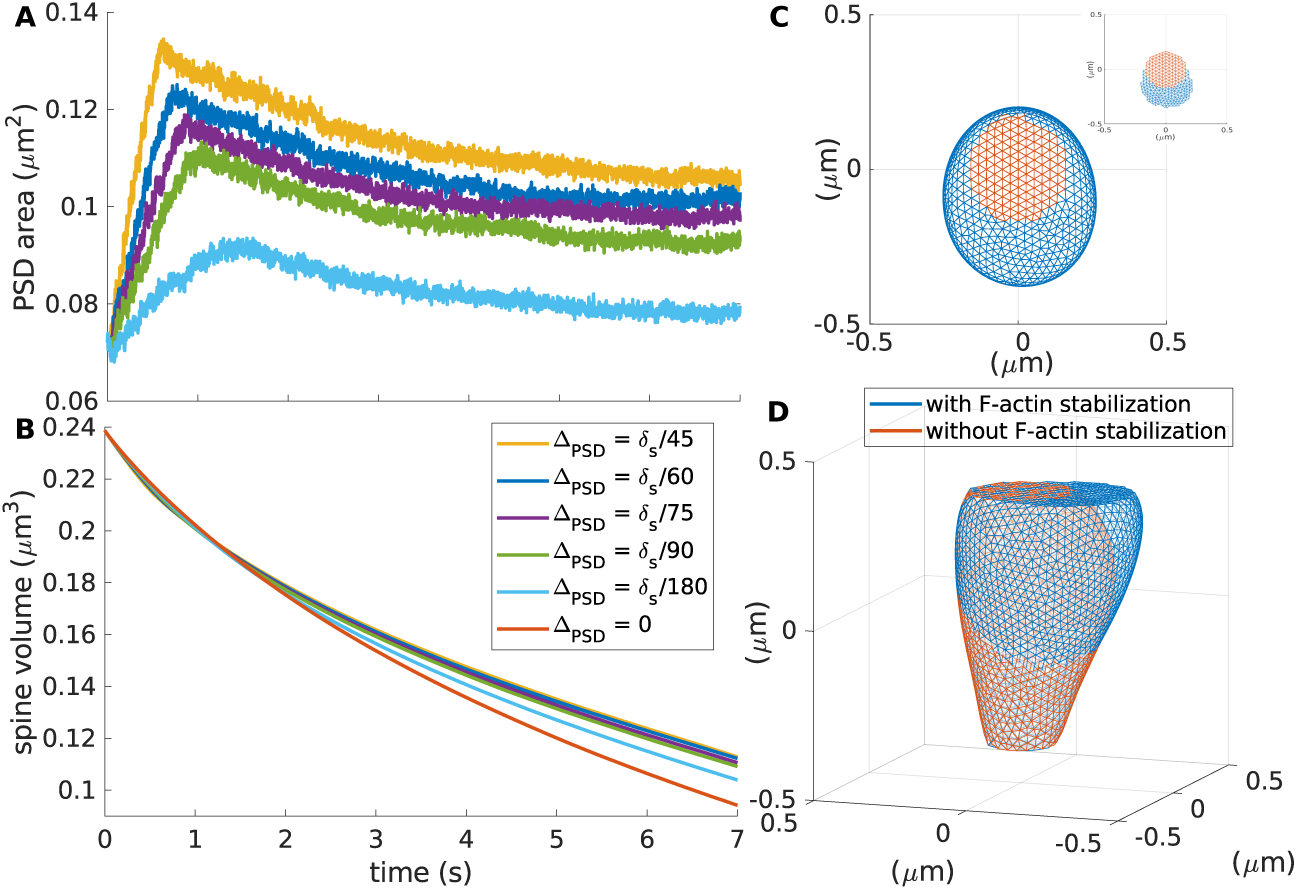
Stabilization of F-actin focus. **A**, PSD surface area over time for different increments of Δ_*PSD*_, color-coded as in B. **B**, spine volume evolution over time for different increments of Δ_*PSD*_. **C**, top view of a spine after 7 minutes of increasing the PSD size (blue, Δ_*PSD*_ = *δ*_*s*_*/*60) and without changing the PSD (orange). Inset shows only the mesh corresponding to the PSD. **D**, front view of the spines in C.

Increasing the PSD area leads to an increase in spine volume, the higher Δ_*P SD*_ the bigger the spine (Fig. 5B). The difference between spines with and without the proposed F-actin stabilization is shown in Figures 5C-D. After seven minutes, the spine without F-actin stabilization (in orange) reduces its initial volume by 60.56% whilst the spine with F-actin stabilization (in blue) decreases by 52.99% with a PSD increase of 42.87%. At this point, the proportion between spine head volume and PSD size, proposed by Arellano *et al*. [16], is kept: PSD size (0.1030 *μ*m) ≈ 0.88 × volume (0.88×0.1122*μ*m^3^ = 0.0987 *μ*m^3^).

Therefore, our simulations show that the F-actin promoting full membrane fusion in an exocytotic event, can be stabilized and serve as anchoring place for PSD proteins. Although the stabilization of this pool of F-actin leads to a asymmetrical growth of the PSD, it does not perturb the shrinkage of the spine. We assumed that the F-actin stabilization occurs at a faster rate than published measures on increase in PSD proteins concentrations [5,9]. Further experiments are needed to test whether this is true in the spine.

## Discussion

We exploited a mathematical model of dendritic spines to test various hypotheses regarding the role and distribution of F-actin polymerization after induction of LTP. Our model operates in a physiological temporal and spatial regime; how-ever, an exact fitting would require additional experimental data, which are currently not available. We adopted a framework where we considered membranes as a two-dimensional elastic continuum [22] without accounting for lipid dynamics [23]. Moreover, we implemented a continuous force generated by actin polymerization rather than simulating the stochastic dynamics of actin [15, 21, 24]. With this framework, we were able to quantify spine enlargement and local membrane tension force. Unlike other 3D models of dendritic spines [14, 25], our model allows for asymmetries that prove to be crucial for membrane properties, such as the force generated by the membrane tension.

Our results indicate that F-actin polymerization increases the force generated by membrane tension and that such an increase is faster for a single focus of polymerization of actin filaments within the spine than for a spine with multiple foci. Moreover, this force is higher if the focus is located near the PSD. Since membrane tension serves as a signal for exocytosis [12], these results are in line with experimental data that show a specialized zone near the PSD for exocytosis of recycling endosomes containing glutamate receptors [17]. These experiments demonstrate that exocytosis is abrupt and massive, and it occurs in an all-or-non fashion in a zone defined by t-SNARE syntaxin 4 [17], but they did not investigate the effects on membrane tension. However, Kliesch *et al*. showed that membrane fusion efficiency mediated by SNARE proteins increases with membrane tension [11] and Grafmüller *et al*. suggested that tension has to increase to start vesicle fusion [26]. Therefore, we propose that t-SNARE syntaxin 4 and an increase in membrane tension, due to actin polymerization, have to co-locate to trigger exocytosis in spines.

Each step of exocytosis is aided by the actin cytoskeleton in different ways [27]. Actin also executes different functions depending on the vesicle size, which dictates the duration of the exocytotic event [27]. Here, instead of modeling all the exocytotic phases, we investigated whether actin polymerization induces a full merge of the Ω-profile. Since exocytosis allows the membrane to achieve spine enlargement [18], this full merge is important. We found that, indeed, surface area and volume increase when actin polymerization is present. Further studies could consider a more detailed description of the force performed by actin filaments, like in previous studies of endocytosis [28].

Our model shows that the F-actin pool, responsible for increasing the spine size, and the force generated by the membrane tension also promotes full merge of the Ω-profile. Therefore, we investigated whether this pool could account for spine stabilization, when it is not longer polymerizing at a fast rate. For this, we assumed that F-actin slowly stabilizes and serves as a scaffold for proteins and receptors at the PSD. We found that without this mechanism, the spine shrinks to its original size. Therefore, the here-proposed F-actin pool, that needs to polymerize at a single location close to the PSD to elevate the membrane tension and trigger exocytosis, acts as the “enlargement pool” proposed by Honkura *et al*. [7]. Furthermore, this actin may act as a synaptic “tag” that captures newly synthesized LTP-related proteins [29, 30].

In conclusion, using our model we were able to investigate in detail the relationship between cell morphology, mechanical properties and LTP-induced events in the dendritic spines. This is a first step towards an in sillico structural model of LTP, that can be extended to include a more accurate description of actin filaments and lipid dynamics. Moreover, further work can link to AMPAR dynamics, as described in [31, 32].

## Methods

### Model

Following our previous work [15], we use a 3D mesh of *n*_*vertices*_ vertices at positions **x**^*k*^ = (*x*^*k*^, *y*^*k*^, *z*^*k*^) ∈ ℝ^3^, *k* ∈ {1, 2, 3, …, *n*_*vertices*_} to represent the spine membrane. At each time-step Δ_*t*_, the vertices move according to

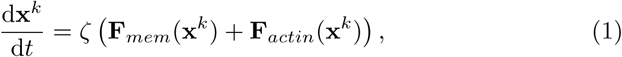

where **F**_*actin*_ is the force generated by F-actin polymerization, given by

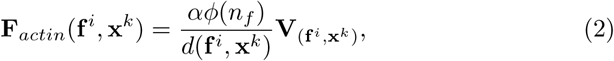

with *d*(**f** ^*i*^, **x**^*j*^) representing the distance between the *i*th actin polymerization focus initial position **f** ^*i*^ and the *k*th mesh vertex **x**^*k*^. We assume that F-actin extends and branches from this location. Here, 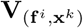 is the normalized direction vector from the focus to the mesh vertex, *α* is the strength of F-actin influence and 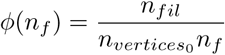 = is a constant proportional to the number of F-actin *n*_*fil*_ observed experimentally [25] and inversely proportional to the number of actin nucleation locations *n*_*f*_ and the number of vertices of the “resting shape” 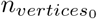 (in black, Fig. 1A).

In Eq. (1), *ζ* is a proportionality constant and **F**_*mem*_ is the force generated by the membrane, given by

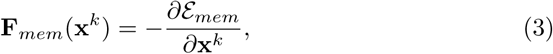

where

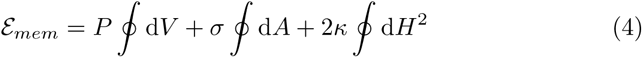

represents the membrane energy. Here, *P* is the difference between internal and external pressure, *V* the volume, *σ* the surface tension, *A* the surface area, *κ* the bending modulus, and *H* the mean curvature. *P, σ* and *κ* are constant parameters that depend on the cell type [33]. The last term of Eq. (4) corresponds to the bending energy, described by Helfrich [34–36], which increases when the membrane is deformed and induces forces that return the membrane to its resting shape. The first and second term in Eq. (4) correspond to the volume and surface area constrains, respectively. See Figure 1A for a sketch of these forces and [15] for details of the calculations.

### Implementation

As in [15], we run the simulations in MATLAB on a desktop computer. At each time-step, Eq. (1) is solved using a classical Runge-Kutta algorithm. Importantly, for numerical accuracy, the mesh has to be isotropic and conserve the number of neighbors of each vertex. Hence, remeshing is needed. For this we use the `remeshing.m` function in [37], that is based on Openmesh [38], with three iterations and a target length of *δ*_*s*_. Parameters for the simulation are given in Table 1.

**Table 1:** Model Parameter Values.

| Symbol | Unit | Definition | Value | Source |
| --- | --- | --- | --- | --- |
| $\Delta t$ | s | length of the time-step | 1/8 | [15] |
| $r_s$ | $\mu\text{m}$ | initial spine radius | 0.4 | To match measures in [16] |
| $\delta_s$ | $\mu\text{m}$ | target length of and edge | 0.03 | [15] |
| $r_{neck}$ | $\mu\text{m}$ | neck radius | 0.0796 | [16] |
| $r_{PSD_0}$ | $\mu\text{m}$ | initial PSD radius | 0.1744 | To match measures in [16] |
| $h_{neck}$ | $\mu\text{m}$ | value for fixing the neck | -0.3920 | Calculated as in [15] |
| $h_{PSD}$ | $\mu\text{m}$ | value for fixing the PSD | 0.36 | Calculated as in [15] |
| $P$ | $\text{pN}\mu\text{m}^{-2}$ | difference between internal and external pressure | 75 | Calculated as in [15] |
| $\sigma$ | $\text{pN}\mu\text{m}^{-1}$ | surface tension | 15 | [15] |
| $\kappa$ | $\text{pN}\mu\text{m}$ | bending modulus | 0.18 | [15] |
| $\alpha$ | pN | strength of filament influence | 3.8 | [15] |
| $\zeta$ | $\mu\text{m}^2\text{s}^{-1}\text{pN}^{-1}$ | strength of force update | 0.004 | [15] |
| $n_{fil}$ | 1 | number of actin filaments in the spine head | 70 | [25] |
| $\delta_s$ | $\mu\text{m}$ | target length of an edge | 0.03 | [15] |
| $\Delta_{PSD}$ | $\mu\text{ms}^{-1}$ | PSD increasing rate | $\delta_s/60$ | - |

The resting shape (black mesh in Fig. 1A) is found as in [15]. In short, from a sphere with radius *r*_*s*_, the points corresponding to the neck and PSD are fixed to *h*_*neck*_ and *h*_*P SD*_, respectively, and **F**_*actin*_ is set to **0**. We let the shape evolve until it settles. In this resting shape, the tracking points are selected to be isotropic with edge length 2*δ*_*s*_. At each time-step, each term of **F**_*mem*_ (Eq. (3)) is calculated for these points. Particularly, for each tracking point **x**^*j*^, the force generated by the membrane tension is given by 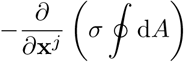. Therefore, our measure of the force generated by tension is equivalent to the mechanical work needed to increase the surface area of the spine. To compare the force generated at different locations, we take its norm ‖ · ‖. The location of the F-actin polymerization foci are selected by either choosing a vertex of the resting shape and scale it 99% or by remeshing a scaled version of the resting shape (99%) with a larger vertex length. This way, the foci are equally distributed within the spine.

For the stabilization of F-actin, at each time-step, the PSD radius increases at a rate of Δ_*PSD*_ and F-actin corresponding to spine enlargement within this radius is stabilized. This is done by fixing the mesh vertices that are within the increased radius in the *x − y* coordinates with ‖*z − h*_*PSD*_‖ ≤ 0.001 to a hight of *h*_*PSD*_ This allows to asymmetrically fix the PSD at the locations where the spine experience growth. Selecting a larger tolerance for the *z* value leads to a more isotropic enlargement of the PSD.

### Code Availability

Custom computer code used to generate the findings of this study will be made available in Github.

## Acknowledgements

We thank the German Science Foundation through the Collaborative Research Center SFB-1286, Project C3 for funding this research.

## Author Contributions

MB-Q performed research and wrote the first draft. MB-Q and FW contributed to the study concept. FW edited the paper.

## Competing Interests Statement

The authors declare no competing interests.

## Notes

### Competing Interest Statement

The authors have declared no competing interest.

