## Supplemental Figure 1 for "Exploring new roles for actin upon LTP induction in dendritic spines"

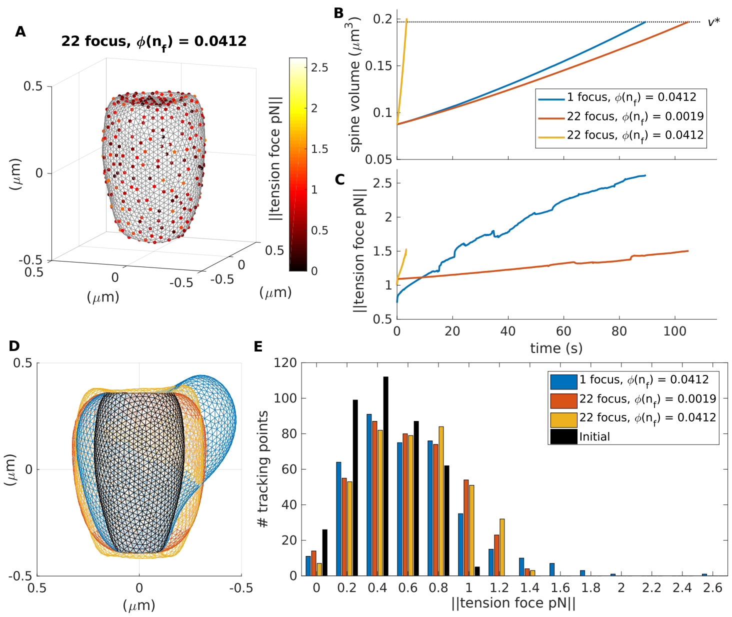


Figure S1: **Spine enlargement upon LTP for different locations of the F-actin polymerization foci and $\phi(n_f)$.** **A**, shape of a spine with 20 equally distributed polymerization foci and $\phi(n_f) = 0.0412$ (gray) at time when it reaches a volume of $v^*$. Dots are the tracking points color-coded for the membrane tension force. **B**, spine volume evolution over time, color-coded for different parameters. Dotted black line denotes $v^*$. **C**, the evolution of the membrane tension for the tracking point with maximum membrane tension when the spine reaches a volume of $v^*$. **D**, spine shapes for different parameters when they reached a volume of $v^*$, color-coded as in B. This plot shows the $y - z$ axis, slid at $x = 0$. Black shape corresponds to the resting shape. **E**, histogram of the distribution of the force generated by membrane tension measured at the tracking points for the spines in D.
